# A neural mass model for the EEG in ischemia

**DOI:** 10.1101/2023.04.07.535995

**Authors:** Manu Kalia, Sophie L.B. Ligtenstein, Hil G.E. Meijer, Michel J.A.M. van Putten

## Abstract

Normal brain function depends on continuous cerebral blood flow for the supply of oxygen and glucose, and is quickly compromised in conditions where the metabolic demand cannot be met. Insufficient cerebral perfusion can result in ischemic stroke, with symptoms ranging from loss of motor or language function to coma, depending on the brain areas affected. Cerebral ischemia also results in changes in the electroencephalogram. Initially, a reduction of the frequency of the rhythms occurs. Depending on the depth and duration of energy deprivation, this eventually leads to the disappearance of all rhythmic activity. Here, we study the relationship between electroencephalogram (EEG) phenomenology and cellular biophysical principles using a model of interacting thalamic and cortical neural masses coupled with energy-dependent synaptic transmission. Our model faithfully reproduces the characteristic EEG phenomenology during acute cerebral ischemia and shows that synaptic arrest occurs before cell swelling and irreversible neuronal depolarization. The early synaptic arrest is attributed to ion homeostatic failure due to dysfunctional Na^+^/K^+^-ATPase. Moreover, we show that the excitatory input from relay cells to the cortex controls rhythmic behavior. In particular, weak relay-interneuron interaction manifests in burst-like EEG behavior immediately prior to synaptic arrest. We corroborate our observations with human EEG data from patients undergoing carotid endarterectomy and patients after cardiac arrest with a postanoxic encephalopathy. The model thus reconciles the implications of stroke on a cellular, synaptic and circuit level and provides a basis for exploring other multi-scale therapeutic interventions.

**Significance statement:** Reliable synaptic transmission and preservation of ion gradients across cellular membranes are essential for physiological brain function and consume significant energy. During cerebral ischemia, synaptic arrest occurs early due to energy deprivation (ED), which is characterized clinically by the loss of physiological electroencephalographic (EEG) rhythms. In this work, we explore connections between cellular and network behavior during ED by means of a novel computational model that describes ion dynamics in the cortex and thalamus, and resulting EEG. We reproduce characteristic EEG behavior during ED and show that synaptic arrest occurs before other pathologies like swelling and depolarization. Moreover, we predict that low excitatory thalamocortical projections cause burst-like EEG patterns before synaptic arrest, which may explain observations regarding post-stroke synaptic reorganization.

## 1 Introduction

Stroke is one of the leading causes of death in the world today [1], with approximately 11% of global deaths, as per the World Health Organisation. Ischemic stroke, in particular, accounts for 87% of all stroke cases and is caused by a blockage in blood supply to the brain. This blockage results in a lack of oxygen and glucose, compromising various energy-dependent processes, such as synaptic transmission and maintenance of membrane potentials [2]. In the core region of the affected area, cerebral blood flow (CBF) is less than ≈ 10 ml/100g/min, which leads to irreversible loss of function occurring within minutes. In the surrounding area, known as the penumbra, CBF is in the range of 10-40 ml/100g/min, and significant neuronal dysfunction exists with potential for recovery, depending on the depth and duration of the remaining perfusion [3]. In the penumbra, synaptic transmission failure is the main cause of loss of function.

Oxygen and glucose are essential for the synthesis of adenosine triphosphate (ATP), which is mainly consumed by the Na^+^/K^+^-ATPase (NKA). The NKA maintains ion homeostasis at the synapse by exchanging 3 Na^+^ ions for 2 K^+^ ions per ATP molecule consumed against their respective concentration gradients. At the synaptic level, the vesicular-ATPase (vATPase) also consumes ATP to maintain the pH level of neurotransmitter-carrying vesicles. The proton gradient is vital to the efficient packing (endocytosis) and release (exocytosis) of these vesicles. Synaptic recycling is interrupted during mild ischemia in the form of limited endocytosis or exocytosis [4]. If energy is restored sufficiently fast, synaptic function recovers completely. However, persistent transmission failure is possible after prolonged ischemia [5]. We note that synaptic transmission failure can occur without changes in membrane potentials or baseline ion concentrations [3].

If the ATP depletion is more severe, resting membrane potentials will also change, as the NKA cannot compensate for the non-zero transmembrane ion currents. The resulting depolarization, in turn, causes further large fluxes of the ions Na^+^, K^+^ and Cl^−^ across the membrane along their concentration gradients, and a net intracellular increase in Na^+^ and Cl^−^ occurs [6]. The increase in intracellular osmolarity will subsequently induce cytotoxic edema [7].

Changes in neuronal function resulting from ischemic stroke, in particular, are also reflected in the electroencephalogram if the cortex is involved (EEG) [8, 9, 10]. In the clinic, EEG monitoring is used in patients at risk for cerebral ischemia, for instance, during carotid endarterectomy [9, 11, 12] or to assess recovery in patients with a postanoxic encephalopathy after cardiac arrest [13, 14]. The EEG changes in acute ischemia are well characterized and depend on the depth and duration of the ischemia. Initially, higher frequencies are suppressed, subsequently followed by the emergence of slower rhythms in the delta range, and finally, all rhythms may disappear [15, 9]. However, the precise biological mechanisms that result in such changes in the EEG are not well known.

In this work, we aim to explore how the breakdown of cellular synaptic function may be observed in EEG using a biophysical approach. For this, we propose a mathematical model of coupled neural masses, where the coupling is based on population-averaged synaptic behavior. Population-averaged ion dynamics of the neural masses result in a firing-rate function dependent on membrane and Nernst potentials. Subsequently, this firing rate is used to compute synaptic currents from one neural mass to another. Each neural mass is surrounded by a finite extracellular bath containing oxygen, and energy deprivation is modeled by transiently depriving the bath of oxygen. We use the model to explain the link between cellular synaptic inhibition and evolution of the EEG rhythms and explore the differential sensitivity of NKA and vATPase function to energy availability.

Neural mass models are widely used to describe population synaptic dynamics based on a population-averaged firing rate [16, 17, 18, 19, 20]. Such models have been successfully applied to explain neuropathologies such as epilepsy [21, 22, 23, 24] and neurodegenerative diseases [25, 26]. Recently, in [27], the Epileptor model [28] was combined with ion dynamics to describe seizure-like events in epilepsy. The Liley model [17] in particular is conductance-based and thus more biophysically interpretable. The model has been extended to produce EEG rhythms associated with postanoxic encephalopathy [10]. Spiking neuron models have also been used to model the effects of stroke on thalamocortical circuits [29]. On a cellular level, ion homeostasis in neurons, astrocytes and the tripartite synapse are modeled using biophysical principles [30, 6, 31, 32, 33]. In some of these works, inhibition of Na^+^/K^+^-ATPase (NKA) is used to model the consequences of energy deprivation.

To our knowledge, there are no computational models that reconcile single-ion and neural mass approaches in the context of stroke. Following work on single-ion dynamics in [6, 33], we present for the first time a biophysical model of a neural mass that depends on ion dynamics of Na^+^, K^+^ and Cl^−^. We obtain dynamics of ion concentrations, membrane potentials, cellular volumes and synaptic currents in one formalism. This approach allows us to extend the idea of bistability observed in previous work [33, 6] to the synaptic context. In particular, we show corroboration with experimental literature by modeling mild ischemia by slow (de)activation of synapses. Moreover, we show that inhibiting the NKA and vATPase results in several phases of synaptic rhythms - from healthy to complete synaptic arrest and cellular depolarization.

We further explore the implication of stroke on neural circuitry in the context of functional reorganization. First, we show that excitatory thalamic relay cells govern normal EEG behavior. Next, for weak relay-interneuron interaction, we find the interface between rhythms and synaptic arrest is characterized by different types of mixed-mode oscillations [34], which are burst-like behaviors that arise from slow changes in cellular function.

We also show that these observations and predictions are consistent with actual clinical EEG data acquired from two patient groups, recorded as part of routine clinical care. This included EEG data from patients undergoing carotid endarterectomy. A standard procedure during this procedure is temporary clamping of the carotid artery to assess if sufficient collateral flow is present; if not, this will be reflected as ipsilateral *α*-slowing and emergence of *δ*-activity. For completeness, slowing of the EEG during test-clamping warrants temporary shunting by the vascular surgeon in order for the procedure to be carried out without risks for permanent ischaemic brain injury [9]. We also present EEG data from a cardiac arrest patient with severe postanoxic encephalopathy. In these patients, the EEG showed synchronous bursts with multiple oscillations. Both observations are reliably reflected in our model simulations by EEG slowing and mixed-mode oscillations, respectively.

The paper is organized as follows: we begin with a section on the model construction and its equations, followed by a brief summary of the data we present to support our predictions. This is followed by results and a discussion of our findings.

## 2 Materials and Methods

The model scheme used in this work is shown in Fig. 1-A. We model two neural regions, cortex and thalamus. The thalamus provides excitatory drive to the cortex and is not subjected to energy deprivation for all simulations shown in this work. For each neural region, we model two local populations, inhibitory and excitatory, that interact via synaptic connections. In the cortex, we model excitatory pyramidal neurons (P) and inhibitory interneurons (I). Similarly, we include excitatory reticular (R) and inhibitory relay (S) neurons for the thalamus. The model for each population is composed of two parts, averaged ion dynamics and the synaptic component. All four local populations are modeled as point neuronal compartments with averaged ion dynamics and embedded in a shared, finite extracellular space. Within these compartments, dynamics of Na^+^, K^+^ and Cl^−^ are modeled via the activity of ion channels and cotransporters. Passive osmotic diffusion enables the compartments to swell or shrink via the movement of water. Each population’s ion dynamics produces a net population firing rate, which activates the AMPA (excitatory) or GABA (inhibitory) channels of other populations, leading to a cascade of excitation and inhibition in the system, see Fig. 1-B.

**Figure 1:**
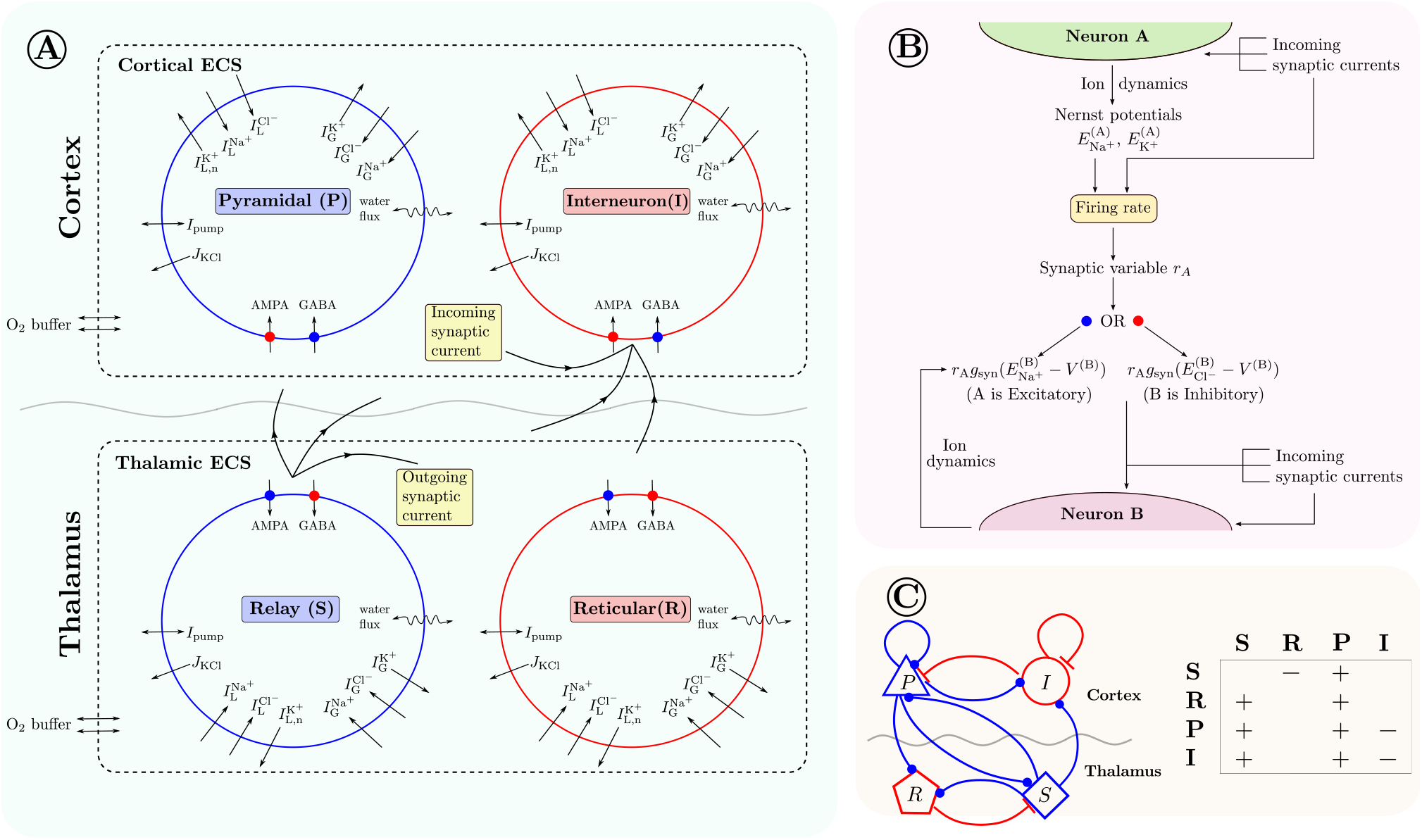
A model of interacting neural populations with biophysical dynamics. (A) The model consists of two neural regions: cortex and thalamus, each composed of two populations. Thus we model four populations, inhibitory interneurons (I), excitatory pyramidal neurons (P), thalamic relay neurons (S) and thalamic reticular neurons (R). The averaged ion dynamics of Na^+^, K^+^ and Cl^−^ for each population are modeled with an averaged single-neuron formalism, where equations are chosen from [6]. Two populations from the same region are enclosed in a finite extracellular space, with an external bath of O_2_. (B) The averaged ion dynamics produce population synaptic currents via a firing rate function which depends on incoming currents and Nernst potentials. (C) The populations are connected via synaptic currents; the network architecture is taken from [35]. Each mark in the table describes an excitatory (plus) or inhibitory (minus) connection from column population to row population.

The single-neuron model from Dijkstra et al. [6] is used to model the averaged ion dynamics of every population. Additionally, the architecture of the neural mass network is chosen from [35]. The EEG signal is then assumed to emerge from the net synaptic current of the pyramidal population.

In the sections ahead, we explain the equations used to describe ion dynamics in a single neural region. The model contains two such neural regions - cortex and thalamus - for which the equations are identical except for parameter values.

### 2.1 Ion dynamics

For each neural region, ion homeostasis in the two populations is modeled based on the averaged behavior of single-neuron ion dynamics. As a result, the fast dynamics corresponding to action potential generation are averaged out, leaving slower dynamics such as ion homeostasis and EEG rhythms intact.

In particular, we describe the dynamics of the molar amounts 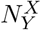 of ions *Y* ∈ *{*Na^+^, K^+^, Cl^−^*}* in populations *X* ∈ *{P, I, S, R}*. The dynamics of population volumes *W*_*X*_ are described by the osmotic imbalance generated by concentrations [*X*]_*Y*_ with respect to the extracellular space. As the total volume and number of ions are kept constant for each neural region, the extracellular volumes and concentrations are given by conservation laws. We fix constants *C*_*Y*_ and *W*_tot_ that refer to the total number of ions of type *Y* and total regional volume, respectively. Thus, the extracellular components are given by,

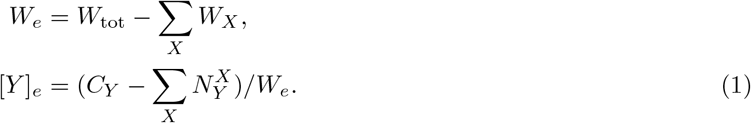

The dynamics of the intracellular components of a population *X* are given by the following differential equations,

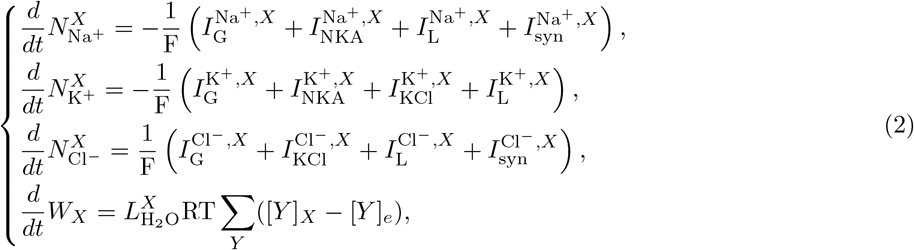

where *F, R* and *T* are Faraday’s constant, the gas constant and fixed temperature, respectively. The subscript *e* indicates an extracellular quantity. 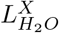 is the average water permeability of the intracellular compartments of population *X*. The various currents 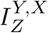 correspond to averaged ion currents of ion *Y* generated by ion channels and cotransporters *Z* in population *X*. Now, we briefly describe each ion current 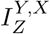. The synaptic currents 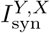 are explained later.

#### Voltage-gated and leak channels

The voltage-gated channels 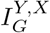 and leak channels 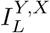 are formulated in the form of Goldman-Hodgkin-Katz (GHK) currents, as done in [6, 33]. The currents are given by,

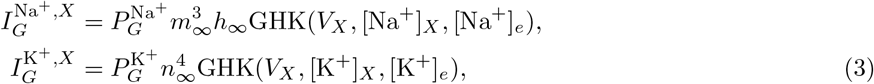

where *m, h* and *n* are gating variables, 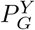 are ion channel permeabilities and *V*_*X*_ is the averaged membrane potential of population *X*. The expression for *V*_*X*_ follows from [6],

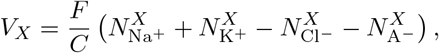

where *C* is the averaged membrane capacitance of population *X* and A^−^ are large impermeant anions in the intracellular space of the population. The function GHK(*V*_*X*_, [*Y*]_*X*_, [*Y*]_*e*_) is given by,

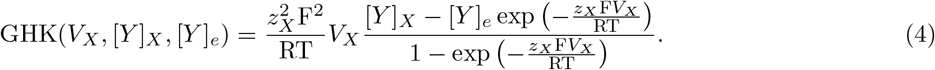

where *z*_*X*_ is the valence of ion species *X*. The terms *m*_∞_, *h*_∞_ and *n*_∞_ are given by,

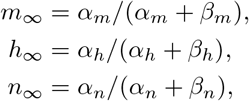

where

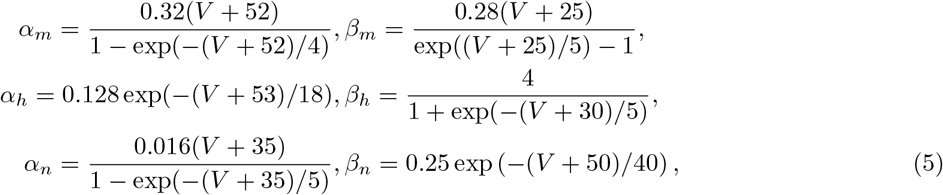

where the subscript was dropped from *V* for convenience. The expression for the voltage-gated Cl^−^ channel follows from [6] and is given by,

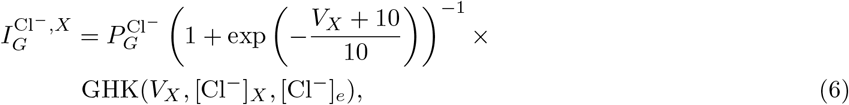

and the leak currents are given by,

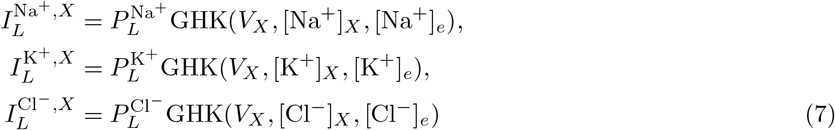

with *P*_*L*_ the leak channel permeabilities.

#### KCl cotransporter

We model the flux as the difference of the K^+^ and Cl^−^ Nernst potentials as in [36],

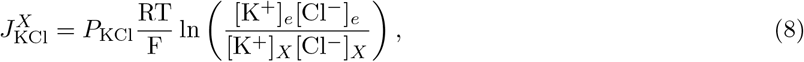

where *P*_KCl_ is the strength of the cotransporter. The corresponding K^+^ and Cl^−^ currents are given by

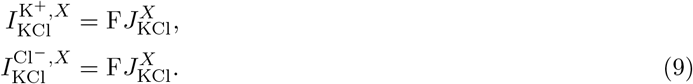

#### Na^+^**/**K^+^**-ATPase (NKA)**

The Na^+^/K^+^-ATPase is modeled as a function of intracellular Na^+^, extracellular K^+^, extracellular Na^+^ and extracellular O_2_. The current follows the model from [37] and is given by

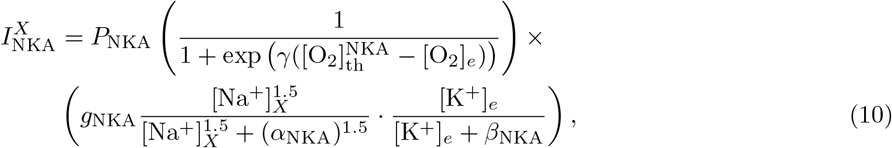

where 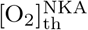 is the oxygen threshold for healthy activity. The functions *g*_NKA_ and *σ* are given by

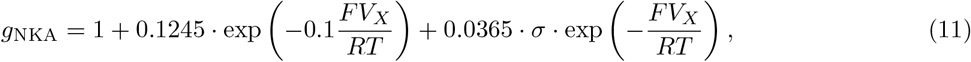

and

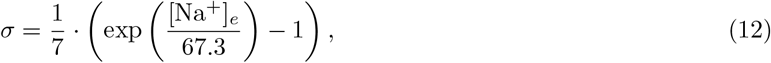

where *P*_NKA_ is the NKA permeability or the pump strength. The corresponding Na^+^ and K^+^ currents are given by

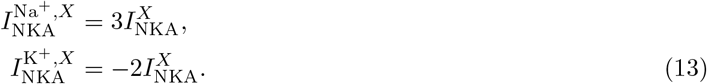

### 2.2 Synaptic dynamics

Next, for the two local populations, we add synaptic currents, which describe the connections to each other and themselves, as shown in Fig. 1. Later on, we extend this to two regions. Synaptic currents depend on a firing rate function, which describes the firing frequency of the population as a function of membrane voltage and magnitude of external current. Usually, this is modeled as a sigmoidal function that depends on a membrane potential threshold. This work proposes a more realistic firing rate function that depends on incoming synaptic currents and Nernst potentials of Na^+^ and K^+^.

#### Biophysical firing rate function

The firing rate function is computed using numerical continuation data of a Hodgkin-Huxley model, which does not have the pump currents, the KCl cotransporter currents and the synaptic currents. From our biophysical setup, the dynamics of the average population membrane potential *V* (subscript dropped for convenience) is given by,

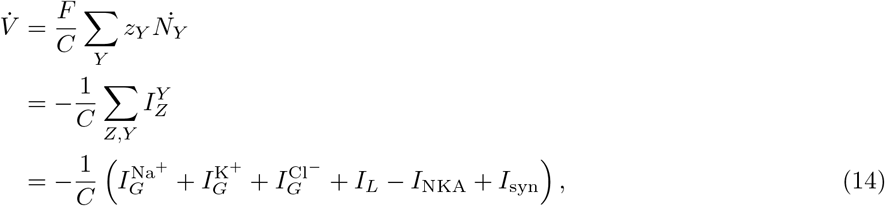

where *I*_*L*_ is the averaged leak current and *I*_syn_ is the total synaptic current

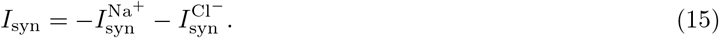

The Hodgkin-Huxley model is given by,

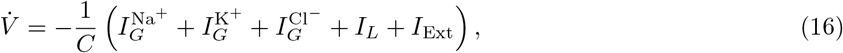

where *I*_Ext_ is external input current. Comparing equations (14)-(16), we see that setting *I*_Ext_ = −*I*_NKA_ + *I*_syn_ in Eq. (16) gives our system Eq. (14). This property is exploited to compute a firing rate function that not only depends on external current, but also on the Nernst potentials of Na^+^ and K^+^. This is done in the following way:

- Perform continuation of the stable periodic orbit in the Hodgkin-Huxley model with respect to three parameters: *E*_K+_, *E*_Na+_ and *I*_Ext_. The existence of the periodic orbit is in a compact domain, marked by two bifurcations on the boundary: a saddle-node of periodic orbits to the right, and a supercritical Hopf to the left, see Fig. 2 (left).
- Next, approximate a smooth function *FR* that describes the frequency of the computed periodic orbits in the previous step, as a function of *E*_K+_, *E*_Na+_ and *I*_Ext_.
- Now, we set the input *I* to the firing rate function as *I*_Ext_ + *I*_syn_ − *I*_NKA_ instead, where *I*_Ext_ is the external current supplied to the population.

**Figure 2:**
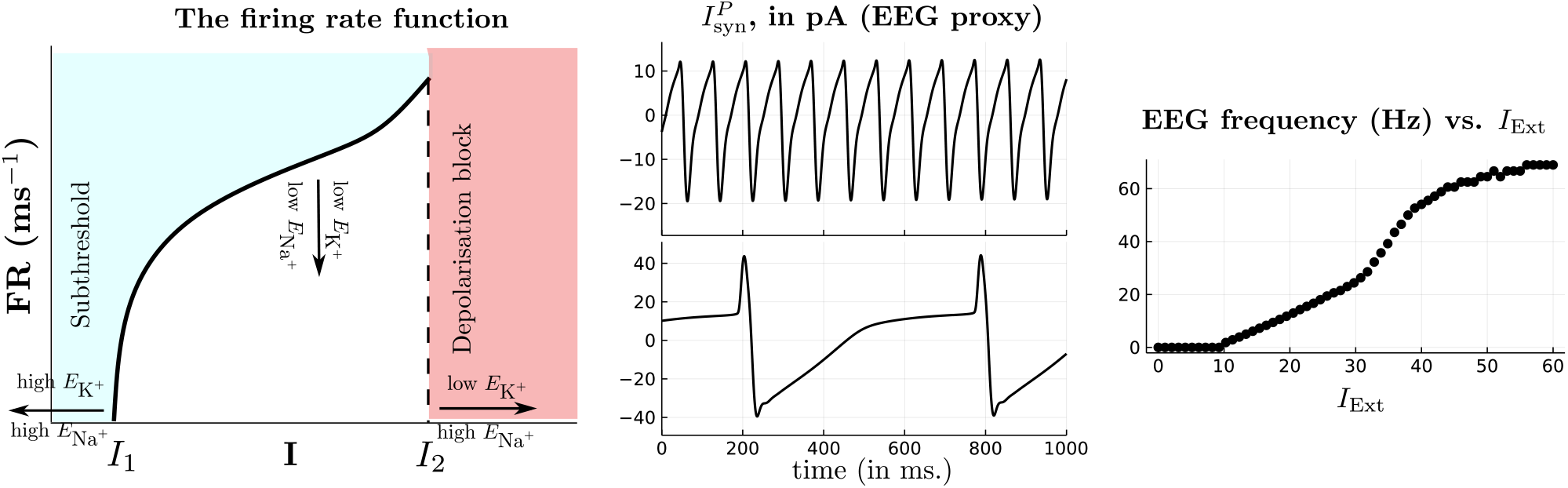
Firing rate and signaling dynamics at baseline. (Left) Plot of the biophysical firing rate function Eq. 17 against input current *I* for fixed *E*_Na+_ and *E*_K+_. The firing rate is characterized by two thresholds of zero firing rate. The first corresponds to subthreshold behavior, and the latter signifies the depolarization block, where averaged population membrane potentials saturate at pathological levels. (Middle) Plots of the net pyramidal synaptic current 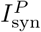 against time under two different conditions. The top trace corresponds to baseline parameters, which give rise to *α* oscillations (≈12 Hz). In the bottom trace, the synaptic constants 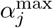 and *β*_*j*_ are reduced to 10% of baseline values which gives rise to *δ* oscillations (≈ 2 Hz). Note the increased amplitude of the *δ* oscillations. (Right) Frequency vs input current to the thalamic relay neurons *I*_Ext_. Increasing input current past a threshold (≈ 10 pA) substantially increases the frequency. For the rest of the simulations, baseline input is set to 20 pA.

The firing rate function *FR* is thus expressed as,

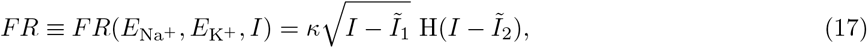

where *I* refers to external input current, H is the standard Heaviside function and 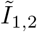 are Nernst potentialdependent thresholds. The functions *κ* and 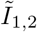 are given by fitted cubic polynomial expansions as follows,

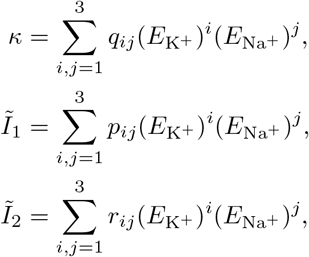

where the fitting constants *q*_*ij*_, *p*_*ij*_ and *r*_*ij*_ are given in Table 5.

#### Synaptic currents and mild ischemia

From the firing rate function *FR*, we compute the synaptic currents 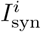 for a population *i*. First, we define the synaptic current 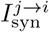 as the current from population *j* to *i*. The net synaptic current for a population *i* is then given by,

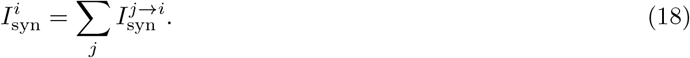

The current 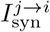 depends on the synaptic gating variable *r*_*j*_ and maximal current of population *i*. The current is modeled as a voltage-gated channel as follows,

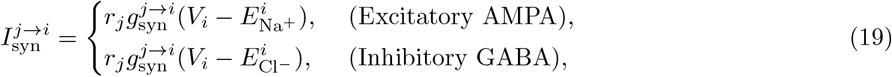

where 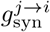 are connectivity constants. They are conductances that serve as network connection parameters. The dynamics of the synaptic variable *r*_*j*_ is given by a typical gating equation and depends on the firing rate *FR*_*j*_ of population *j*,

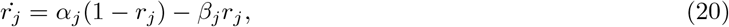

where

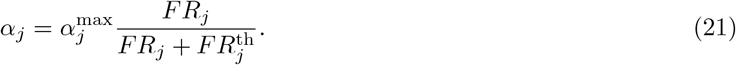

The threshold 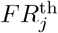 depends on the properties of the synaptic behavior of population *j*. For inhibitory populations, this threshold is kept higher than for excitatory populations. The currents 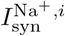 and 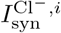 are then given by,

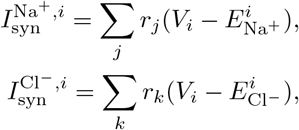

where *j* and *k* span over excitatory (E,S) and inhibitory (I,R) populations, respectively. The proxy for EEG is computed as the pyramidal synaptic current 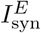 that is run through a band-pass filter, filtering the signal to only show rhythms in the 0.1-40 Hz range.

### 2.3 Oxygen dynamics and simulating energy deprivation

We explicitly model oxygen dynamics as a way of performing transient energy deprivation. First, an infinitely large bath of oxygen is supplied to each neural region. This bath contains a constant concentration of oxygen [O_2_]_bath_ = 2 mM. The diffusion of oxygen in the extracellular space happens via linear diffusion, where the baseline value is set at [O_2_]_*e*_ = 1.75 mM, as per [38]. The influx of oxygen in the extracellular space is balanced by the consumption of NKA. The dynamics of [O_2_]_*e*_ are thus given by [38],

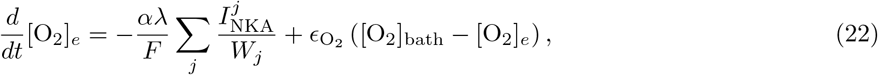

where *j* runs over all populations from a single neural region. The constants *α* and *λ* are O_2_ consumption rates from [38] and the constant 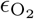 is the O_2_ diffusion constant. Ischemia is simulated by transiently reducing the constant [O_2_]_bath_ to lower values before bringing it back to baseline, as done in [38]. For each population, we obtain four equations describing ion dynamics, given by Eq. (2) and the dynamics of its synaptic variable *r*, given by Eq. (20). Next, for each neural region, we obtain two equations for extracellular O_2_ dynamics, given by Eq. (22). In this work, we model four populations of two neural regions, so in total, we have 22 differential equations for our working model.

### 2.4 Parameters and simulation

All the parameters used in the model equations are shown in Tables 1-5. Most of the parameters are chosen directly from previously published work, except the ones shown in tables 4 and 1. The parameters shown in Table 1 are estimated from baseline conditions by fixing state variables to their baseline values and setting the right-hand side of system equations to zero. The synaptic parameters, shown in Table 4, are empirically estimated such that *α* rhythms are produced following 20 pA external input current.

**Table 1:** Parameters estimated from baseline conditions. Units are presented in the same manner as they are implemented in the Julia code.

| Constant | Value | Description |
| --- | --- | --- |
| $P_L^{\text{Na}^+}$ | $1.28 \times 10^{-6} [1000 \mu\text{m}^3(\text{ms})^{-1}]$ | $\text{Na}^+$ leak channel permeability. |
| $P_L^{\text{K}^+}$ | $1.252 \times 10^{-5} [1000 \mu\text{m}^3(\text{ms})^{-1}]$ | $\text{K}^+$ leak channel permeability. |
| $P_L^{\text{Cl}^-}$ | $2.812 \times 10^{-6} [1000 \mu\text{m}^3(\text{ms})^{-1}]$ | $\text{Cl}^-$ leak channel permeability. |
| $W_e$ | 16 $[1000 \mu\text{m}^3]$ | Baseline extracellular volume. |
| $N_{A-}^X$ | 302.0105 [fmol] | Amount of impermeant anions in a population soma. |
| $N_{A-}^e$ | 416.10 [fmol] | Amount of impermeant anions in the extracellular space. |
| $C_{\text{Na}^+}$ | 2484 [fmol] | Total amount of $\text{Na}^+$ ions in a neural region. |
| $C_{\text{K}^+}$ | 628 [fmol] | Total amount of $\text{K}^+$ ions in a neural region. |
| $C_{\text{Cl}^-}$ | 2192 [fmol] | Total amount of $\text{Cl}^-$ ions in a neural region. |
| $W_{\text{tot}}$ | 20 $[1000 \mu\text{m}^3]$ | Total volume of a neural region. |
| $\epsilon_{\text{O}_2}$ | $1.1175 \times 10^{-4} [\text{ms}^{-1}]$ | Oxygen diffusion constant |
| $g_{\text{vATP}}$ | 0.2409 [pA] | maximum vATPase current |

**Table 2:** Initial values for the various states in the model in all neural regions and populations *X*. These values correspond to ‘baseline’ conditions, and are used to estimate unknown parameters. Units are presented in the same manner as they are implemented in the Julia code.

| Constant | Value | Description |
| --- | --- | --- |
| $[\text{Na}^+]_X$ | 13 [mM] | Taken from [70]. |
| $[\text{K}^+]_X$ | 145 [mM] | Taken from [6]. |
| $[\text{Cl}^-]_X$ | 7 [mM] | Taken from [6]. |
| $W_X$ | 2 $[1000 \mu\text{m}^3]$ | Baseline neuronal soma volume (taken from [6]) |
| $r_X$ | 0 | Initial synaptic variable |
| $[\text{O}_2]_e$ | 1.25 [mM] | Baseline oxygen concentration in the extracellular space [38] |

**Table 3:** Parameters associated with ion dynamics. Units are presented in the same manner as they are implemented in the Julia code.

| Constant | Value | Description |
| --- | --- | --- |
| C | 20 pF | Membrane capacitance [6] |
| F | 96485.333 $[\text{C mol}^{-1}]$ | Faraday’s constant |
| R | 8314.4598 $[\text{C}(\text{mV})(\text{mol K})^{-1}]$ | Universal gas constant |
| T | 310 K | Room temperature [6] |
| $P_G^{\text{Na}^+}$ | $8 \times 10^{-4} [1000 \mu\text{m}^3(\text{ms})^{-1}]$ | Voltage-gated $\text{Na}^+$ channel permeability [6] |
| $P_G^{\text{K}^+}$ | $4 \times 10^{-4} [1000 \mu\text{m}^3(\text{ms})^{-1}]$ | Voltage-gated $\text{K}^+$ channel permeability [6] |
| $P_G^{\text{Cl}^-}$ | $1.95 \times 10^{-5} [1000 \mu\text{m}^3(\text{ms})^{-1}]$ | Voltage-gated $\text{Cl}^-$ channel permeability [6] |
| $\alpha_{\text{NKA}}$ | 13 [mM] | NKA: Half-saturation concentration for intracellular $\text{Na}^+$ [33] |
| $\beta_{\text{NKA}}$ | 0.2 [mM] | NKA: Half-saturation concentration for extracellular $\text{K}^+$ [33] |
| $P_{\text{KCl}}$ | $1.3 \times 10^{-6} [\text{fmol} (\text{ms mV})^{-1}]$ | KCl cotransporter strength [6] |
| $L_{H_2O}^X$ | $2 \times 10^{-14} [1000 \mu\text{m}^3 \text{mPa}^{-1} \text{ms}^{-1}]$ | Neuronal membrane water permeability [6] |
| $\alpha$ | 0.1666 | Conversion factor [38] |
| $\lambda$ | 1 | Relative cell density [38] |
| $[\text{O}_2]_{\text{bath}}$ | 1.75 [mM] | Baseline bath oxygen concentration [38] |
| $\gamma$ | 20 $[\text{mM}^{-1}]$ | Sigmoidal slope constant |
| $[\text{O}_2]_{\text{th}}^Y$ | Varies, see Fig. 3 [mM] | Oxygen threshold for ATPase |

**Table 4:** Parameters associated with synaptic dynamics. Parameters are empirically adjusted. Units are presented in the same manner as they are implemented in the Julia code.

| Constant | Value | Description |
| --- | --- | --- |
| $g^{S \rightarrow R}$ | 0.3 [nS] | Population connectivity |
| $g^{S \rightarrow E}$ | 0.5 [nS] | Population connectivity |
| $g^{S \rightarrow I}$ | 0.2 [nS] | Population connectivity |
| $g^{R \rightarrow S}$ | 2 [nS] | Population connectivity |
| $g^{E \rightarrow S}$ | 0.5 [nS] | Population connectivity |
| $g^{E \rightarrow R}$ | 0.1 [nS] | Population connectivity |
| $g^{E \rightarrow E}$ | 0.3 [nS] | Population connectivity |
| $g^{E \rightarrow I}$ | 0.5 [nS] | Population connectivity |
| $g^{I \rightarrow E}$ | 5 [nS] | Population connectivity |
| $g^{I \rightarrow I}$ | 2.5 [nS] | Population connectivity |
| $FR_S^{\text{th}}, FR_E^{\text{th}}$ | 0.2 [ms<sup>-1</sup>] | Excitatory firing rate threshold |
| $FR_R^{\text{th}}, FR_I^{\text{th}}$ | 0.5 [ms<sup>-1</sup>] | Inhibitory firing rate threshold |
| $\alpha_E$ | 12.5 [ms<sup>-1</sup>] | Pyramidal synaptic activation constant |
| $\alpha_I$ | 5 [ms<sup>-1</sup>] | Interneuron synaptic activation constant |
| $\alpha_S$ | 1.25 [ms<sup>-1</sup>] | Relay synaptic activation constant |
| $\alpha_R$ | 0.5 [ms<sup>-1</sup>] | Reticular nuclei synaptic activation constant |
| $\beta_E$ | 3 [ms<sup>-1</sup>] | Pyramidal synaptic deactivation constant |
| $\beta_I$ | 0.03 [ms<sup>-1</sup>] | Interneuron synaptic deactivation constant |
| $\beta_S$ | 0.3 [ms<sup>-1</sup>] | Relay synaptic deactivation constant |
| $\beta_R$ | 0.003 [ms<sup>-1</sup>] | Reticular nuclei synaptic deactivation constant |

**Table 5:** Coefficients of polynomials 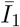, *κ* and 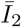.

| Coefficient ($\bar{I}_1$) | Value | Coefficient ($\kappa$) | Value | Coefficient ($\bar{I}_2$) | Value |
| --- | --- | --- | --- | --- | --- |
| p00 | -213.8 | q00 | 0.287 | r00 | -185.9 |
| p10 | -8.404 | q10 | 0.01094 | r10 | -4.877 |
| p01 | -3.007 | q01 | 0.001027 | r01 | -3.674 |
| p20 | -0.1111 | q20 | 0.0001692 | r20 | -0.002785 |
| p11 | -0.07862 | q11 | -8.42e-6 | r11 | -0.1044 |
| p02 | -0.001213 | q02 | -4.839e-5 | r02 | 0.01561 |
| p30 | -0.0005002 | q30 | 8.167e-7 | r30 | -1.17e-5 |
| p21 | -0.0005221 | q21 | -3.156e-7 | r21 | 1.897e-5 |
| p12 | -1.483e-6 | q12 | -4.421e-7 | r12 | 0.0006927 |
| p03 | 3.779e-5 | q03 | 2.764e-7 | r03 | 0.0001089 |

The parameters *g*^*i*→*j*^ are determined using the neural mass network architecture from [39, 35]. These parameters have to be adjusted from the original sources where they are fluxes (in units volt second) as in this work, they represent ion channel conductances (in units siemens). The parameters *α*_*i*_ and *β*_*i*_ are fixed in accordance with three assumptions. First, the deactivation constant *β*_*i*_ for inhibitory populations is kept at least a factor 10 lower than others to accommodate for the slowly deactivating GABA_*B*_ receptor [40]. Second, the parameters *g*^*j*→*i*^ for inhibitory populations *j* are kept a factor 10 higher than the rest, as the resting Cl^−^ reversal potential is close to the resting membrane potential resulting in a small net GABA current. Finally, the parameters *α*_*i*_ and *β*_*i*_ are set to be a factor 10 lower than their cortical counterparts to ensure slow-wave input from the thalamus to cortex.

All simulations ahead are performed in Julia, and the code is available online at: https://github.com/mkalia94/BioNeuralMass. For continuation methods, we use Matcont [41].

#### Baseline behavior

We use the ion-based neural mass model to predict EEG behavior during mild ischemia and the effect of network heterogeneity on rhythmic behavior. The model is simulated by supplying the relay cell with an external input current of 20 pA. This stimulus excites the system and generates a robust *α* rhythm, which we define as baseline activity. This is shown in Fig. 2 (middle, top).

The input current is chosen such that the model shows an *α* rhythm at baseline. In Fig. 2 (right), we plot resulting EEG frequencies against input current *I*_Ext_, which shows a steady increase of frequencies after a threshold (≈ 10 pA). Our choice corresponds to an EEG frequency of ≈ 13 Hz.

### 2.5 EEG recording and clinical data

To illustrate the biological realism of our model, we compare model simulations with human EEG recordings. For all EEG recordings, electrodes were applied according to the international 10/20 system, using 19 channels. Electrode impedances were kept below 5 kΩ. Sampling frequency was set to 256 Hz. A Neurocenter EEG system (Clinical Science Systems, the Netherlands) was used with a TMS-i full band EEG amplifier (TMS international, the Netherlands). We included patients who underwent carotid endarterectomy, a procedure during which temporary clamping of the carotid artery is performed to assess if EEG changes occur that would warrant temporary shunting [9]. In addition, we included a comatose patient after cardiac arrest treated in the intensive care unit. These patients suffered from global cerebral ischaemia, and, depending on the depth and duration of the hypoxic burden, various characteristic EEG patterns can be observed, including abnormal rhythmic patterns, reflecting the severity of the postanoxic damage [42, 13, 43, 44]. All data were fully anonymized before analysis. The Medical Ethical Committee waived the need for informed consent as data collection was part of routine clinical care.

## 3 Results

We use the neural mass model to predict EEG behavior during mild ischemia. Baseline behavior is defined by a steady *α* rhythm. We then subject it to mild ischemia by transiently reducing Na^+^/K^+^-ATPase activity. Further, we explore network heterogeneity by varying parameters 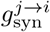.

### 3.1 Delta rhythms emerge from slow (de)activation of synapses

Physiological EEG rhythms in the cortex are characterized by a dominant frequency in the *α* range (8-13 Hz). As a consequence of acute ischemia, these faster rhythms lose power, and slower (*δ*) rhythms emerge [9]. We demonstrate this behavior in Fig. 2. Here, the relay (excitatory thalamic) population is subjected to 20 pA of current, which provides an excitatory drive to the cortical populations. The synaptic currents of all populations then show synchronous rhythmic activity in the *α* range (≈12 Hz). In Fig. 2 (middle, top), we plot the resulting synaptic current from the pyramidal cell (cortex, excitatory), which is used as a proxy for measuring EEG activity. Throughout this simulation, the membrane potentials and ion concentrations do not deviate from baseline conditions.

Next, we perform the same simulation but with mild ischemia. For this setting, we slow the synaptic activation and deactivation constants of the cortical populations, setting them to 10% of their baseline values. As before, we also subject the excitatory thalamic population to 20 pA constant current. The resulting simulation shows rhythmic behavior too, but much slower, i.e., in the *δ* range, see Fig. 2 (middle, bottom).

In our simulations, slowing the synaptic activation and deactivation constants of the cortex results in a *α* to *δ* transition, typical of mild ischemia. Based on this observation, we model the effect of the vesicular ATPase on synaptic dynamics, by changing Eq. 20 to

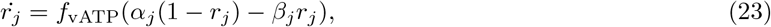

where *f*_vATP_ depends on extracellular O_2_ and models the effect of the vesicular ATPase on the activation and deactivation of the synaptic variable. In the absence of O_2_, it simulates mild ischemia by slowing down gating behavior. The term *f*_vATP_ is modeled as a sigmoid as follows,

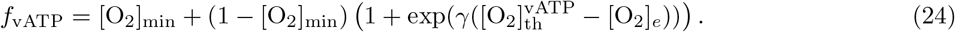

The constant [O_2_]_min_ prevents synapses from completely slowing, and is set to 0.1, or 10% of baseline values. We assume that the vATPase consumes extracellular O_2_ in the same way as the NKA. Thus, Eq. 22 becomes,

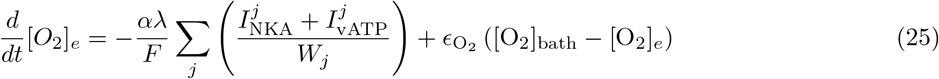

where *I*_vATP_ is the vATPase current associated with oxygen consumption, and is given by,

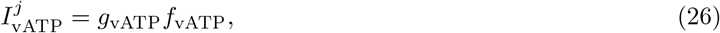

where the maximum current *g*_vATP_ converts the flux to a current and is computed from rest extracellular space happens g baseline conditions. The slowing of *α*-rhythms and the simultaneous emergence of *δ*-rhythms is also evident in patient data collected during carotid endarterectomy, see Fig. 4. We plot the hemisphere-averaged power spectrum of the EEG of seven patients before, during, and after carotid artery clamping. There is a marked slowing of the EEG during the clamping phase (typically starting within 30-60 s), along with an increase in *δ*-rhythms. In all cases, the EEG recovers to baseline *α*-rhythms post-clamping as the blood has now been restored.

### 3.2 Mild ischemia leads to synaptic arrest via multiple phases

Next, we consider our extended model with the vATPase. We simulate energy deprivation in this case by transiently reducing [O_2_]_bath_. We show the result of such simulations in Fig. 3.

**Figure 3:**
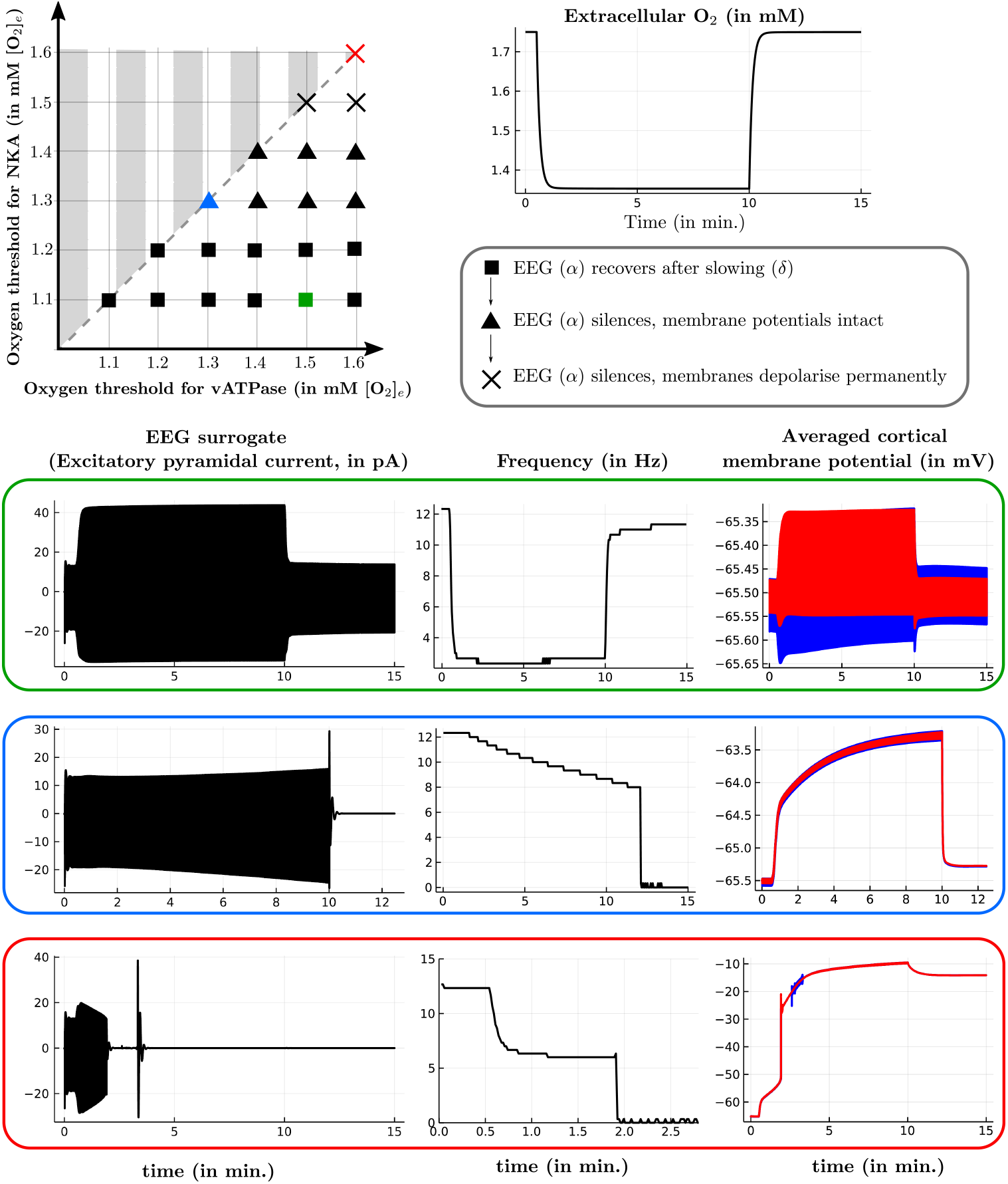
Two-parameter sweep of the model with oxygen dynamics. (Top, left) Two parameter plot of 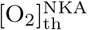 vs 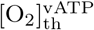 for several values between 1.1 and 1.6 mM. For each parameter pair, the oxygen bath supply [O_2_]_bath_ is reduced to 80% of its original capacity between minutes 0.5 and 10 while simultaneously stimulating the thalamic relay neuronal population with 20 pA for the whole duration of the simulation. This results in the extracellular oxygen trace (Top, right). For each parameter pair, we show the three possible cases (square, triangle, cross) corresponding to the transitional behavior of the EEG. The cases are ordered in pathological effect, and for each case, we plot the corresponding EEG signal behavior (Bottom, left solid black line). The green panel also shows that the *α*-*δ* transition is accompanied by increased amplitude during slowwave activity. Alongside each EEG plot, we also show its frequency-time plot (Bottom, right), which shows the dominant frequency characteristics of the transitional behavior, and the corresponding averaged cortical membrane potentials. The red and blue traces correspond to interneurons and pyramidal cells, respectively.

**Figure 4:**
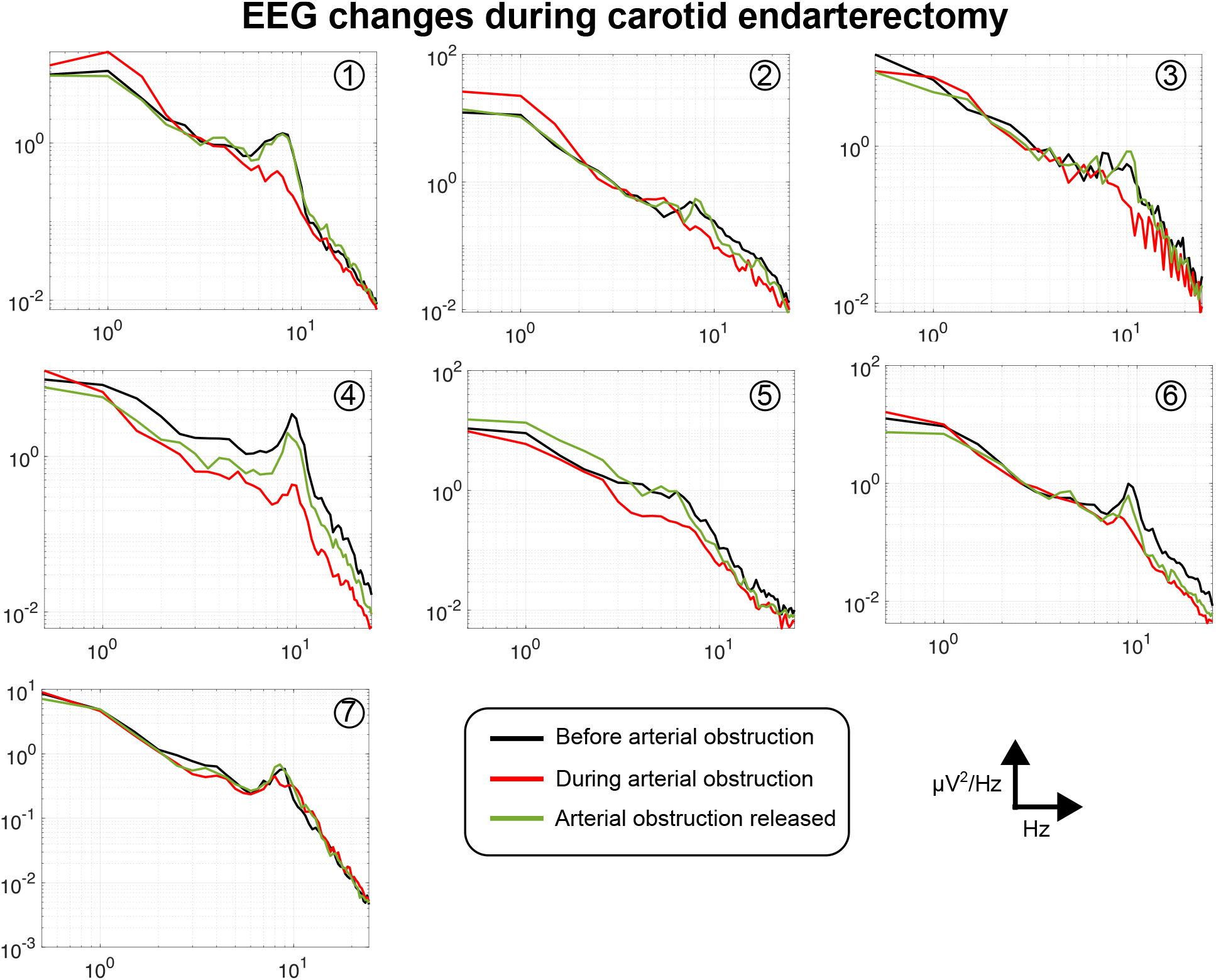
*α* − *δ* transition in patients during temporary carotid artery occlusion. Averaged timefrequency plots of the power spectrum density (PSD) of the EEG obtained from 7 patients during carotid endarterectomy where the carotid artery is temporarily occluded to assess if shunting is needed [9]. Each line corresponds to the PSD of the EEG that is averaged over all channels from a particular hemisphere (left/right). For all patients except 3 and 5, data from the right hemisphere is plotted. For each patient, the PSD is plotted before (black), during (red) and after (green) carotid artery occlusion. The PSD is computed for each signal of length 20 s. Typical duration of temporary clamping is 1-2 minutes. In all patients, a dominant peak frequency in the alpha range (9-10 Hz) is present. Note the reduction or near disappearance of the peak frequency (approximately 9-10 Hz) during arterial occlusion in all patients shown (only in patient 7 this is very limited). Further, in patients 1 and 2, an increase in the delta power (0.5-3 Hz) is observed.

The extracellular oxygen concentration, [O_2_]_bath_, is reduced transiently to 80% of its baseline value between minutes 0.5 and 10. Meanwhile, 20 pA excitatory thalamic stimulation is provided for the full length of the simulation, as before. In Fig 3-top,right we plot the bath oxygen concentration [O_2_]_e_, which shows a characteristic transient profile of rapid initial decrease followed by slow recovery, as seen in previous work [38, 45].

Although the threshold 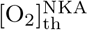 is available in literature [38], the choice for the vATPase threshold 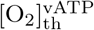is not clear. Given the high ATP demand of the NKA compared to endo/exocytosis [3], we assume that 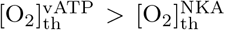. Thus we perform a parameter sweep by varying the two thresholds for various values between [1.1, 1.6]. For each parameter pair, we simulate energy deprivation as before while simultaneously providing excitatory thalamic input throughout the simulation. This setup allows us to observe transitions between several rhythmic patterns. The results of the parameter sweep are shown in Fig. 3-top, left. We observe three distinct behaviors of the EEG signal, and we plot an example for each case. Alongside the EEG, we plot a time-frequency plot of the signal, computed via a moving window for every time point. The first (square) corresponds to a recovery of the *α* rhythm after oxygen restoration. In this case, the EEG may slow down to the *δ* range during low-oxygen conditions. The presence of low-frequency rhythms is also accompanied by a significant increase in EEG amplitude. Recovery occurs for relatively low NKA thresholds.

Upon increasing 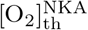slightly, the EEG no longer recovers to the *α* range post oxygen restoration and remains flat (triangle and cross). In this scenario, two subcases are possible. First (triangle), the EEG stays flat after energy restoration, but membrane potentials and ion dynamics recover to baseline. The second (cross) case corresponds to a total loss of synaptic communication and ion homeostasis - the EEG stays flat post energy restoration, and the averaged neuronal membranes are permanently depolarized.

Moreover, varying 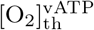 while keeping 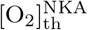 fixed does not change the qualitative behavior of the EEG signal. Thus it suffices to keep these parameters identical.

### 3.3 Relay cells control cortical rhythmic behavior

The global onset of cortical stroke may also have consequences on the thalamocortical circuit itself [46, 47, 48]. Our model contains four thalamocortical connections. These amount to the following parameters being nonzero: 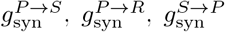 and 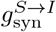. We now vary these parameters to see the effect of severing thalamocortical contact on baseline signaling. First, we are able to immediately neglect two parameters: 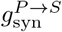 and 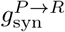. These correspond to the excitatory input provided by cortical pyramidal cells to both thalamic populations. The parameter 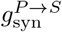 modulates the frequency of baseline behavior but does not drastically affect the signaling process. Further, setting the parameter 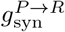 to zero results in no change in qualitative behavior. Thus, we are left with two parameters 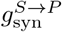 and 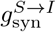, which control the excitatory thalamic input to the cortical populations. We perform a parameter sweep of these parameters in the two-parameter domain, see Fig. 5. Baseline *α* rhythms correspond to the parameter pair (0.5, 0.2).

**Figure 5:**
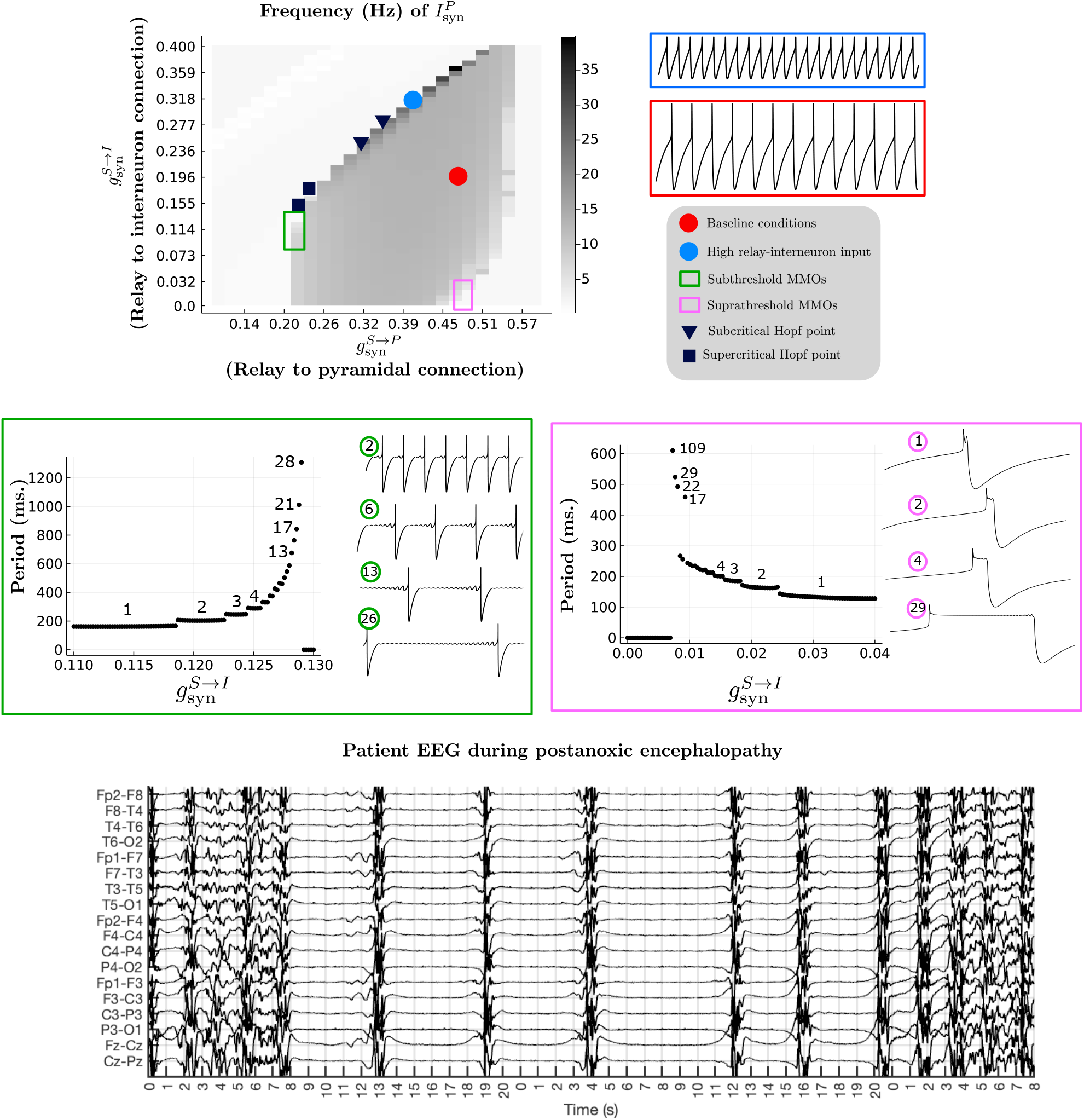
The impact of relay neurons on cortical rhythms. (Left, top) Plot of a two-parameter sweep with parameters 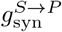 and 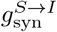, corresponding to relay input to pyramidal neuronal and interneuronal populations, respectively. For each parameter value, the frequency of the corresponding signal 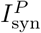 is shown. For a few cases, the corresponding 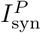 is shown in pA (black trace). Two simulations of regular rhythms are shown for baseline conditions (red) and the near-Hopf case (blue).(Middle) The two insets explore transitions involving mixed-mode oscillations (MMOs) (green and pink). In the green box, the period of 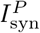 is plotted (black dots) against parameter 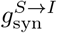 while 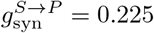, showing a period-adding sequence. The number on the black dots corresponds to the number of subthreshold small amplitude oscillations (SAO). The onset of MMOs coincides with subcritical Hopf bifurcations nearby (black square), which transition to supercritical Hopf bifurcations (black inverted triangle) as 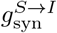 is increased. Similar plots are shown in the pink box, where 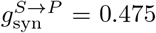. Here, the numbers correspond to the number of suprathreshold SAOs. (Bottom) Plot of the EEG of a patient after a cardiac arrest with severe postanoxic encephalopathy. The EEG shows characteristic bursts [43], similar to the MMOs in model simulations that represent the transition point between healthy and unhealthy EEG behavior.

For each parameter, the resulting frequency of the total pyramidal synaptic current 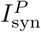 is plotted. For a few cases, corresponding traces of 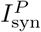are shown. Several interesting behaviors emerge. We observe that rhythms exist for a small window of 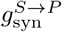. While keeping 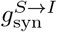 fixed, we see that there is a small window where rhythmic behavior exists. Increasing 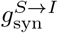 squeezes this window, till there are no more rhythms possible. Baseline *α* rhythms (red point) are structurally stable in a local neighborhood and transit to silent behavior via three thresholds. For relatively low values of the parameter 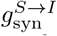 (less than 50% of baseline), two thresholds correspond to sequences of period-adding bifurcations (green and pink box). In both cases, the period-adding sequences are characterized by mixed-mode oscillations (MMOs) [49, 34], see traces in the green and pink box. For low values of 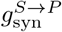 (green box), the MMOs contain subthreshold small-amplitude oscillations (SAOs). For each jump in the period, a new subthreshold SAO is added to the signal. For higher values of 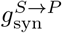 (pink box), the sequence of MMOs contains suprathreshold SAOs. The last threshold (blue) corresponds to a supercritical Hopf bifurcation. Here, regular rhythms disappear while the frequency settles to a fixed high frequency and the amplitude decays.

The onset of MMOs coincides with nearby subcritical Hopf bifurcations, two examples of which are shown in the parameter sweep (black squares). Here, the silent EEG state loses stability without the emergence of stable periodic behavior. Upon increasing 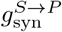, the Hopf bifurcations transition into a supercritical type (black inverted triangles). A small perturbation of the parameter 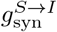 at such a point results in the emergence of stable rhythmic behavior, with low amplitude. The Hopf points were computed via MatCont [41] using one-parameter continuation. It is natural to expect that these points form a continuous curve, where the transition from the supercritical to subcritical case is characterized by a generalized Hopf bifurcation. Due to the slow-fast and multi-scale nature of the system, continuation as a Hopf bifurcation curve was not feasible. However, upon detection of a Hopf point, MatCont also determines whether the Hopf bifurcation is sub- or super-critical. Our one-parameter continuations support the existence of a codim-two bifurcation.

The MMOs with long phases of SAOs are reflected very well in the EEG of a patient suffering from postanoxic encephalopathy. The patient in particular is in a comatose state - on the verge of EEG silencing - which justifies our comparison. The bursts, however, have multiple oscillations interspersed with silence, while the MMOs from our model show a single large oscillation followed by a long period of small amplitude oscillations.

## 4 Discussion

In this work, we have introduced a neural mass model to describe the pathophysiology of mild ischemia in the context of clinical stroke. The model is derived from biophysical principles and encapsulates the dynamical description of four populations in the cortex and thalamus.

### 4.1 Sequence of pathologies in low-oxygen conditions

Mild oxygen deprivation results in a smooth transition from *α* rhythms to the emergence of *δ* activity. In acute hemispheric strokes, the amount of asymmetry in *δ*-power is associated with the neurological status [9, 11, 8, 50, 51, 29]. The appearance of delta activity in acute ischemia is reversible if cerebral blood flow is restored sufficiently fast [9], but after 5-7 minutes of insufficient perfusion, irreversible neuronal damage can occur and EEG changes persist [52]. This is also reflected in the EEG recordings obtained during carotid endarterectomy, see Fig. 4. Temporary clamping of the carotid artery during the procedure can induce energy deprivation, mainly in the ipsilateral hemisphere (the side of the shunting) of the brain, depending on the collateral circulation. If the ipsilateral flow is insufficient for the energy demand, this will quickly result, typically within 30-60 s, in slowing of the EEG [53] and occurrence of hemispheric spectral asymmetries [11, 9]. Here, we average the EEG power spectrum over the ipsilateral hemisphere to demonstrate slowing.

Synaptic transmission is metabolically demanding. At the presynaptic side, ATP is used on several types of ATPase, including the sodium/potassium pump, the Na^+^/Ca^2+^ exchanger, the calcium-ATPase and the vesicular H^+^-ATPase. Postsynaptically, ATP use is larger and primarily needed for restoring ion fluxes involved in synaptic currents [54]. The vATPase is an ATP-driven pump that ensures the necessary proton gradient across the vesicle membrane to allow efficient neurotransmitter secretion in the synaptic cleft [55]. Severe ischemia can result in excitotoxicity, characterized by neurotransmitter accumulation in the cleft and saturation of postsynaptic activity. Mild ischemia mainly affects vesicular endo- and exocytosis [4, 56], resulting in *presynaptic* transmission failure [5, 3].

We capture this phenomenon by modeling the vATPase by a sigmoidal factor that depends on extracellular oxygen. During low oxygen conditions, the vATPase slows down synaptic constants 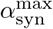 and *β*_syn_, thereby slowing resulting synaptic variables and EEG rhythms, see Figure 5. We found that varying vATPase affinity to available oxygen does not change the sequence of pathological behaviors associated with oxygen deprivation. Upon further reducing energy, EEG rhythms can disappear and remain absent even after energy restoration. Here, ion homeostasis in the population can either stay normal or settle at pathological behavior, see figure* 5. We observe that EEG silencing accompanied by normal ion homeostasis always precedes the case with depolarized populations, upon varying available oxygen or varying NKA/vATP oxygen thresholds. This is attributed to the failure of synaptic rhythmic generation in the model, which precedes failure in ion homeostasis, concurrent with experimental literature on *in vivo* and *in vitro* models [57, 5, 3, 58].

### 4.2 Thalamic relay neurons control baseline activity

Our model contains four connections between the thalamus and cortex. We find that altering thalamocortical connections (connections from thalamic relay cells to interneurons and pyramidal cells) shows burst-like EEG oscillations at the interface between normal rhythms and silent EEG behavior. Cortical ischemic stroke not only causes damage to cortical neurons, but may also result in substantial loss of thalamocortical projection and generally dampens excitability in thalamocortical circuits [59, 46, 60, 61, 62, 44]. This behavior is replicated in the model. For low values of 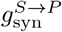, rhythmic activity is lost, see Fig. 5. Moreover, we hypothesize that rhythmic activity exists for a window of relay-pyramidal cell connections. The interface is marked either by fast, low-amplitude oscillations or burst-like behavior.

Mixed-mode oscillations characterize the burst-like behavior for weak relay-interneuron connectivities. MMOs are widely studied dynamical objects that arise in many computational neuroscience models [63, 64, 65, 66, 67, 68]. We observe MMOs of subthreshold and suprathreshold type as seen in previous literature [69], that occur at the border between normal and silent EEG activity. The suprathreshold MMOs are remarkably similar to generalized periodic discharge (GPD) behavior in EEGs, which are hypothesized to result from weak excitatory input to inhibitory interneurons [42, 13], which agrees with our model predictions, see Fig. 5. Moreover, GPDs are also hypothesized to be prevalent at the interface between normal and low-voltage EEG behavior [42]. These events may also be related to pathological burst suppression patterns, which are associated with poor neurological outcome following stroke [43, 10].

We support our prediction with the EEG recorded in a post-cardiac arrest, comatose, patient with a severe postanoxic encephalopathy, showing a characteristic burst-suppression pattern. The observation is consistent with our prediction that bursts are at the interface between complete synaptic arrest (EEG silencing) and normal EEG behavior. The EEG shows multiple oscillations in the burst, while our model simulations only account for a single large oscillation with several small-amplitude oscillations in the ‘silent’ phase. Nevertheless, as there are multiple period-adding windows in the model, we expect more such regimes to exist for other combinations of the synaptic connectivity parameters, where perhaps MMOs with multiple large oscillations exist.

### 4.3 Model assumptions and limitations

Our model has several limitations. For instance, astrocytes are not included in the current model, while their role in ion homeostasis is undisputed [33, 7]. We also assume that ischemia may affect thalamocortical projections, while preserving intrinsic neuronal function in the thalamus. While this is in agreement with reports on postmortem histopathology in patients with a postanoxic encephalopathy after cardiac arrest, showing that cortical neurons can be selectively affected [44], the involvement of the thalamus cannot be excluded. Several model parameters are empirically optimized to ensure baseline *α* activity. However, these choices - including network architecture - are not unique and their perturbation results in no qualitative change in the results shown. Despite these limitations, our simulations show similar EEG characteristics as clinically observed and provide candidate biophysical pathophysiological mechanisms for its generation.

### 4.4 Conclusion

In sum, we constructed a neural mass model based on biophysical principles of ion dynamics, which allows us to make predictions simultaneously about synaptic behavior and ion homeostasis. Our model makes several predictions for EEG rhythms following adapted neuronal activity due to mild ischemia or during post-stroke functional reorganization. High amplitude *δ* rhythms emerge during mild ischemic connections, and bursting behavior via mixed-mode oscillations may manifest as a result of altering thalamocortical connections. The detailed behavior of the model makes it generalizable to other pathological behaviors such as epilepsy, and can be used to investigate network-based and neuronal pathologies simultaneously.

## 5 Appendix

## Notes

https://github.com/mkalia94/BioNeuralMass

### Competing Interest Statement

The authors have declared no competing interest.

